# Repurposed kinase inhibitors and β-lactams as a novel therapy for antibiotic resistant bacteria

**DOI:** 10.1101/199422

**Authors:** Nathan Wlodarchak, Nathan Teachout, Rebecca Procknow, Jeff Beczkiewicz, Adam Schaenzer, Kenneth Satyshur, Martin Pavelka, Bill Zuercher, Dave Drewry, John-Demian Sauer, Rob Striker

**Author notes:** To whom correspondence should be addressed: Rob Striker, Department of Medicine, University of Wisconsin-Madison, 3301 Microbial Sciences Building, 1550 Linden Dr., Madison,WI 53706.

## Abstract

Antibiotic resistant bacteria are an increasing global problem, and pathogenic actinomycetes and firmicutes are particularly challenging obstacles. These pathogens share several eukaryotic-like kinases that present antibiotic development opportunities. We used computational modelling to identify human kinase inhibitors that could be repurposed towards bacteria as part of a novel combination therapy. The computational model suggested a family of inhibitors, the imidazopyridine aminofurazans (IPAs), bind PknB with high affinity. We found that these inhibitors biochemically inhibit PknB, with potency roughly following the predicted models. A novel x-ray structure confirmed that the inhibitors bind as predicted and made favorable protein contacts with the target. These inhibitors also have antimicrobial activity towards Mycobacteria and Nocardia, and normally ineffective β-lactams can potentiate IPAs to more efficiently inhibit growth of these pathogens. Collectively, our data show that *in silico* modeling can be used as a tool to discover promising drug leads, and the inhibitors we discovered can synergize with clinically relevant antibiotics to restore their efficacy against bacteria with limited treatment options.

## Introduction

Antibiotic resistant bacteria are one of the greatest modern health threats and, if left unchecked, will outpace cancer as a cause of death by 2050(O'Neill, 2014). Despite renewed calls for novel approaches to antibiotic discovery and development, new antibiotic discovery continues to lag, and is unprofitable to pharmaceutical companies(Boucher, Talbot et al., 2009). *Mycobacterium tuberculosis* (*Mtb*) is a particular concern as it currently infects two billion people worldwide, and infections with multidrug resistant tuberculosis (MDR) and extensively drug resistant tuberculosis (XDR) are steadily increasing worldwide(WHO, 2015). In the US, 10% of *Mtb* cases are resistant to common treatments, prompting the CDC to list *Mtb* as a pathogen of serious concern(CDC, 2013). Several other related Gram+ pathogens are becoming increasingly difficult to treat in certain settings. *Nocardia spp*. are susceptible to a narrow range of drugs, many of which are not be tolerated in all patients, decreasing second-line options(Schlaberg, Fisher et al., 2014). Most recently, the NIH has listed non-tubercular mycobacterial infections as an area of particular interest due to rapid increases in these infections with similar resistance concerns(NIAID, March 7, 2017).

Mycobacteria antibiotic resistance is multifactorial, including inactivating enzymes and an impenetrable cell wall(Smith, Wolff et al., 2013). Mycobacteria encode β-lactamases and aminoglycoside acetyltransferases specifically restricting the use of the corresponding antibiotics, though some utility remains(Sotgiu, D'Ambrosio et al., 2016, Wivagg, Bhattacharyya et al., 2014). Additionally, mycobacteria readily develop genetic and epigenetic resistance through selection(Smith et al., 2013) and as such, treatment options for mycobacteria have remained limited and stagnant for over thirty years(Cole, 2016). Co-therapy β-lactam antibiotics and β-lactamase inhibitors have shown some success(Diacon, van der Merwe et al., 2016, Ramon-Garcia, Gonzalez Del Rio et al., 2016, Veziris, Truffot et al., 2011), however, new approaches and new targets are essential to diversify treatment options.

Although bacterial signal transduction pathways are not targets of current antibiotics (Coates, Halls et al., 2011), they may represent novel opportunities for drug development. Bacterial eukaryotic-like serine/threonine protein kinases (STPKs) are one example due to their function in regulating transcription, metabolism, cell cycle regulation, virulence and drug resistance(Wright & Ulijasz, 2014). A subset of these, the Penicillin-binding-protein And Ser/Thr kinase-Associated (PASTA) kinases are integral membrane proteins that bind peptidoglycan fragments and regulate growth, cell wall maintenance, metabolism, biofilm formation, and β-lactam resistance(Beltramini, Mukhopadhyay et al., 2009, Liu, Fan et al., 2011, O'Hare, Duran et al., 2008, Shah, Laaberki et al., 2008, Turapov, Loraine et al., 2015). The mycobacterial PASTA kinase, PknB, is essential for cell survival and virulence, making it an ideal drug target(Fernandez, Saint-Joanis et al., 2006). Despite a clear rationale for PknB inhibitors, compounds with biochemical activity thus far have demonstrated poor microbiologic activity(Chapman, Bouloc et al., 2012, Lougheed, Osborne et al., 2011, Naqvi, Malasoni et al., 2014). Pharmacologic inhibition or genetic deletion of PASTA kinases in *Listeria monocytogenes* and MRSA sensitizes these bacteria to β-lactam antibiotics, suggesting this strategy may be useful in other pathogenic PASTA-kinase containing bacteria(Pensinger, Aliota et al., 2014, Vornhagen, Burnside et al., 2015).

Most antibiotic drug development is aimed at targets with no eukaryotic analogs and thus limit the need for selectivity. Selectivity is attainable for structurally conserved targets, and, for example, it was recently shown that mycobacterial proteasome proteins could be selectively targeted despite close structural similar human targets(Totaro, Barthelme et al., 2017), proving that antibiotic development need not be limited to targets with no human homologs. Additionally, human kinase inhibitors are among the most successful drugs of the 21st century, despite kinases having high structural conservation(Cohen, 2002). The structural similarity of PASTA kinases to human STPKs suggests that not only could similar inhibitors be developed, but inhibitors developed for human kinases may be viable lead compounds for bacterial kinases(Pereira, Goss et al., 2011). Interest in developing human kinase inhibitors has resulted in a wealth of both successful and failed compounds(Drewry, Willson et al., 2014, Heerding, Rhodes et al., 2008), many of which have well developed structure-activity relationship (SAR) data, known toxicity profiles, and are synthetically feasible(Wang, Zorn et al., 2014). Thousands of known kinase inhibitor structures are available in digital databases which can be used for computational modelling(Irwin, Sterling et al., 2012, Kim, Thiessen et al., 2016). Re-purposing drugs for other therapeutic targets is common in noninfectious disease research, but has not been applied to antibiotic discovery because most current bacterial targets were chosen because they do not have structural similarity to human proteins. This approach is attractive because it is potentially cost effective and efficient(Fischbach & Walsh, 2009).

Although several groups have modeled inhibitors in PknB, these studies have generally investigated known inhibitors to determine preferred molecular interactions(Naqvi et al., 2014, Punkvang, Hannongbua et al., 2016, Seal, Yogeeswari et al., 2013). Here, human kinase inhibitors were systematically studied *in silico* to identify new PknB inhibitors. A family of our hits was then validated biochemically, structurally and microbiologically. We show that computational methods for bacterial kinase drug discovery can produce valuable active hits. Additionally, for the first time, we combined successful inhibitors with β-lactam antibiotics to treat pathogenic mycobacteria and Nocardia resulting not only in a new class of antibiotics but also a novel co-therapy.

## Results

### The imidazopyridine aminofurazans (IPAs) are predicted to bind PknB with high affinity

We docked over 1000 kinase inhibitors and found that several were predicted to bind PknB with high affinity (**Fig. 1A**). The mean binding energy for all docked ligands was −8.87 kcal/mol with a standard deviation (σ) of 1.25 kcal/mol and a range of −13.01 to −4.29 kcal/mol. We focused on 22 lead compounds which docked in the 95^th^ percentile (2σ). Nine of the 22 compounds fell into three structurally related groups and the rest were dissimilar. Several of the dissimilar compounds are non-specific inhibitors (rapamycin) or are large hydrophobic compounds unlikely to enter cells (zotarolimus). Two of these groups contained compounds predicted to have low solubility and were larger than 500 Da (Lipinski rule violators). This reduced our list to three compounds: GSK2181306A, GSK1007102B, and GSK554170A. GSK1007102B and GSK554170A are structurally similar and were selected for several reasons. The predicted binding mode suggested they made several protein-ligand interactions (hydrogen bonds, stacking interactions, salt bridges), with major interactions along the protein backbone or with conserved residues, making resistance development by single point mutations less likely. They have several hydrogen bond donors and acceptors and are not excessively hydrophobic, making them more likely to enter the cell and easier to formulate(Lipinski, Lombardo et al., 1997, Rouse, Seefeld et al., 2009). These compounds are members of the imidazopyridine aminofurazan (IPA) family which have well-established SAR. We obtained limited quantities of certain compounds through our collaboration(Heerding et al., 2008, Rouse et al., 2009). Additionally, a member of this family, GSK690693, was used in phase I clinical trials with only moderate side-effects, suggesting that it may be a viable starting point for further development(Carol, Morton et al., 2010).

**Figure 1.**
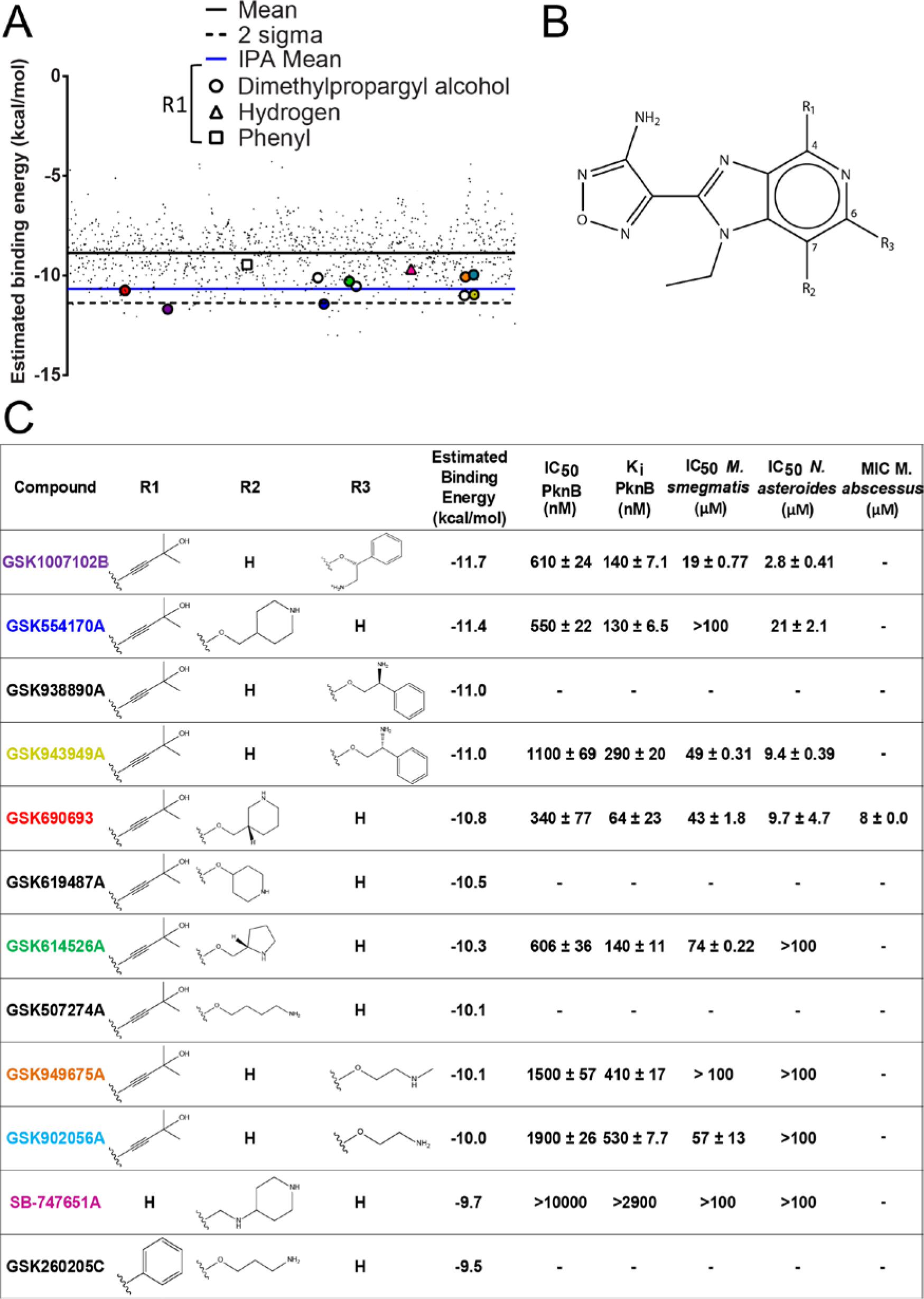
Computationally predicted binding energy, biological IC_50_, and biochemical inhibition trend together for the IPA family of compounds. **A)** A scatter plot showing binding energy values for the *in silico* screen of kinase inhibitors in PknB. Virtual kinase inhibitor libraries were docked in PknB (1O6Y) using Autodock4. Each point represents one compound with the corresponding estimated binding energy plotted along the Y-axis. Points are separated along the X-axis in alphanumeric order. Lower binding energies indicate more favorable binding to the target. The solid black line indicates the mean energy for all three libraries, with the dashed line representing two standard deviations above the mean. The blue line represents the average binding energy of the IPA family (open symbols). Values presented are the lowest energy of 30 trials. Compounds with colored circles were those used in biochemical and microbiological studies, and colors correspond to those in subsequent figures. **B)** The main scaffold structure of the IPAs with relevant carbons and R groups numbered. **C)** Library IPA compounds and related IA compounds are listed along with R group structures, predicted binding energy, biochemical IC_50_ values, calculated K_i_ values, and microbiologic IC_50_ values (drug + β-lactam). GSK690693 (red) was used for crystallographic studies. K_i_ values were calculated using **Equation 1**. Biochemical inhibition curves are shown in **Figure S2**. *Msmeg* and *Nast* inhibition curves are shown in **Figures S6 and S9** respectively. The MIC given for Mab is that of ampicillin at 50μM GSK690693 (**Table S3**).

The IPAs consist of an imidazopyridine core with an aminofurazan ring attached to the imido group (Fig. 1B). The core is modified at three different R groups on carbons 4 (R_1_), 6 (R_2_) or 7 (R_3_) with the R_2_ and R_3_ positions of known IPAs being mutually exclusive: if one has a functional group attached, the other will be a hydrogen only(Heerding et al., 2008, Rouse et al., 2009). The R_1_ position was optimized for Akt as a dimethyl propargyl alcohol, whereas cell entry was modulated with diverse R_2_/R_3_ groups(Heerding et al., 2008). The IPAs docked in PknB with a range of −11.7 to −9.5 kcal/mol, with more cyclic R_2_/R_3_ groups demonstrating higher affinity than those with linear groups (Fig. 1C), suggesting a potential SAR.

### IPAs are potent biochemical inhibitors of PknB and require a functional group at R_1_ for activity

To test whether our modeling predicted biochemical activity, we first characterized PknB kinetics using GarA, which is phosphorylated on T22 by PknB(O'Hare et al., 2008). Kinetic parameters for PknB were previously estimated at sub-saturating conditions using a construct lacking the juxtamembrane domain(Villarino, Duran et al., 2005), however autophosphorylation of residues in this domain are required for full activation of the kinase(Lombana, Echols et al., 2010). These residues are autophosphorylated during expression in *E. coli*, therefore observed *in vitro* kinase activity is on T22 of GarA only(Duran, Villarino et al., 2005). We characterized the enzyme kinetics under a matrix of conditions where both substrates were saturating in order to determine accurate kinetic parameters(Marangoni, 2003). Although kinases vary widely in their enzymatic properties, our values (**Table S1**) were comparable to several other eukaryotic S/T kinases(Brignola, Lackey et al., 2002, Marangoni, 2003, Zhang, Zhang et al., 2006). Interestingly, when the data were plotted with respect to GarA, the allosteric sigmoidal model was preferred (**Fig. S1**). PknB dimerizes both *in vitro* and *in vivo*, making it plausible that allosteric cooperativity is occurring(Barthe, Mukamolova et al., 2010, Lombana et al., 2010)and the calculated Hill coefficient in our model was 1.9, suggesting that at least two subunits are interacting(Marangoni, 2003).

Using drug concentrations from 0-10μM (**Fig. S2**), K_i_ values ranged from 64nM to 530nM (Fig. 1C). K_i_ values for GSK1007102B, GSK554170A, GSK690693, and GSK614526A were not statistically different, but were significantly better than GSK943949A, GSK949675A, and GSK902056A. SB-747651A had no inhibitory activity at concentrations tested, suggesting the importance occupying the back pocket. Inhibitors with cyclic R_2_/R_3_ groups varied statistically from linear R_2_/R_3_ groups (GSK902056A and GSK949657A) by a p <0.01 (Fig. 2). Inhibition of substrate phosphorylation in a dose dependent manner was confirmed using a radiometric assay (**Fig. S3**). Collectively, these results demonstrate that our *in silico* model accurately predicted inhibitors with nanomolar biochemical activity.

**Figure 2.**
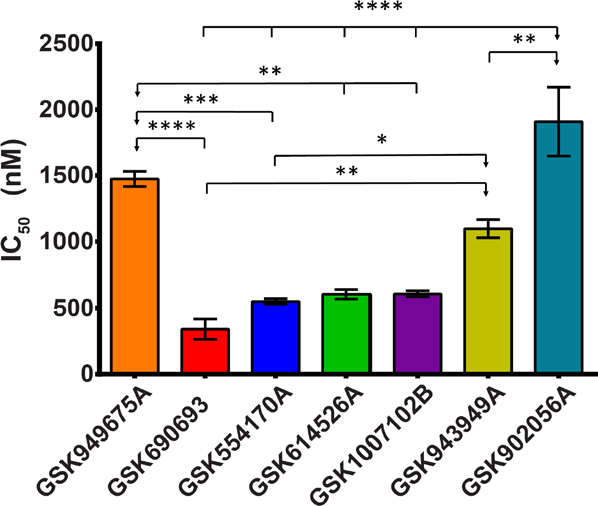
IPAs have significantly different PknB inhibition. IC_50_ values are shown for each compound along with the SEM. One way ANOVA was performed and statistically different groupings are denoted by ticks with arrows indicating the compound from which the group differs. Statistical significance is given by p < 0.05 (*), p < 0.01 (**), p < 0.001 (***), and p < 0.0001 (****). Statistics were calculated using PRISM (GraphPad), and the Tukey correction was used for multiple comparisons. IC_50_ and K_i_ values are given in **Figure 1**, and compound colors correspond to those in **Figure 1**.

### GSK690693 co-crystallizes with PknB, binding in an ATP-competitive manner with several critical interactions

Several PknB structures were previously elucidated, but none with a specific kinase inhibitor bound to the active site(Lombana et al., 2010, Ortiz-Lombardia, Pompeo et al., 2003, Wehenkel, Fernandez et al., 2006, Young, Delagoutte et al., 2003). Additionally, several PknB inhibitors were previously modeled *in silico*, but structural confirmation of these models was not established(Chapman et al., 2012, Lougheed et al., 2011, Naqvi et al., 2014, Seal et al., 2013). To confirm our model and facilitate future drug development insight, we co-crystallized the catalytic domain of PknB with GSK690693, a compound biochemically equivalent to our best drugs and commercially available in large quantities. PknB-GSK690693 crystals grew in 14 different screening conditions (0.9% hit rate). Upon refinement, well-formed crystals grew fully in 2-5 days. Crystals diffracted to 2.0Å and the model was refined at 2.2Å with an R/R_free_ of 19.89/23.26% and favorable geometric parameters (**Table S2**). The overall structure is a bi-lobed globular protein 59.2Å long and 39.5Å across (Fig. 3A). The backbone of the model was almost identical to other published structures at similar resolution (1O6Y (2.2 Å) and 2FUM (2.9 Å)) and similar to the Stk1 structure (4EQM (3.0 Å)) from *Staphylococcus aureus*(Ortiz-Lombardia et al., 2003, Rakette, Donat et al., 2012, Wehenkel et al., 2006). The activation loop (residues 164-177) was not visible in the structure. The initial F_o_-F_c_ map after molecular replacement indicated a well outlined area of unmodelled density in the ATP binding pocket of the active site, extending into the pocket behind the catalytic site (Fig. 3B). GSK690693 satisfied the unmodelled density and bound in the same orientation as it binds in Akt2 (3D0E), except for the R_2_ piperidine group which extends into solution rather than interacting with the ATP binding floor (**Fig. S4**)(Heerding et al., 2008). The aminofurazan ring makes two hydrogen bonds to the backbone of the hinge region of PknB (E93 and V95), and the dimethylpropargyl alcohol makes two hydrogen bonds to the back pocket via the backbone of F157 and sidechain of E59. The piperidine ring makes a weak hydrogen bond to the backbone of the P-loop (F19) through an ordered water. The catalytic lysine (K40) forms a hydrogen bond (2.9Å) with the nitrogen of the pyridine ring (Fig. 3C & D). M145 and M155 make stacking interactions with the imidazopyridine core, and M92 makes a stacking interaction with the aminofurazan ring (Fig. 3D). The ligand is also coordinated through 8 hydrophobic interactions (Fig. 3C).

**Figure 3.**
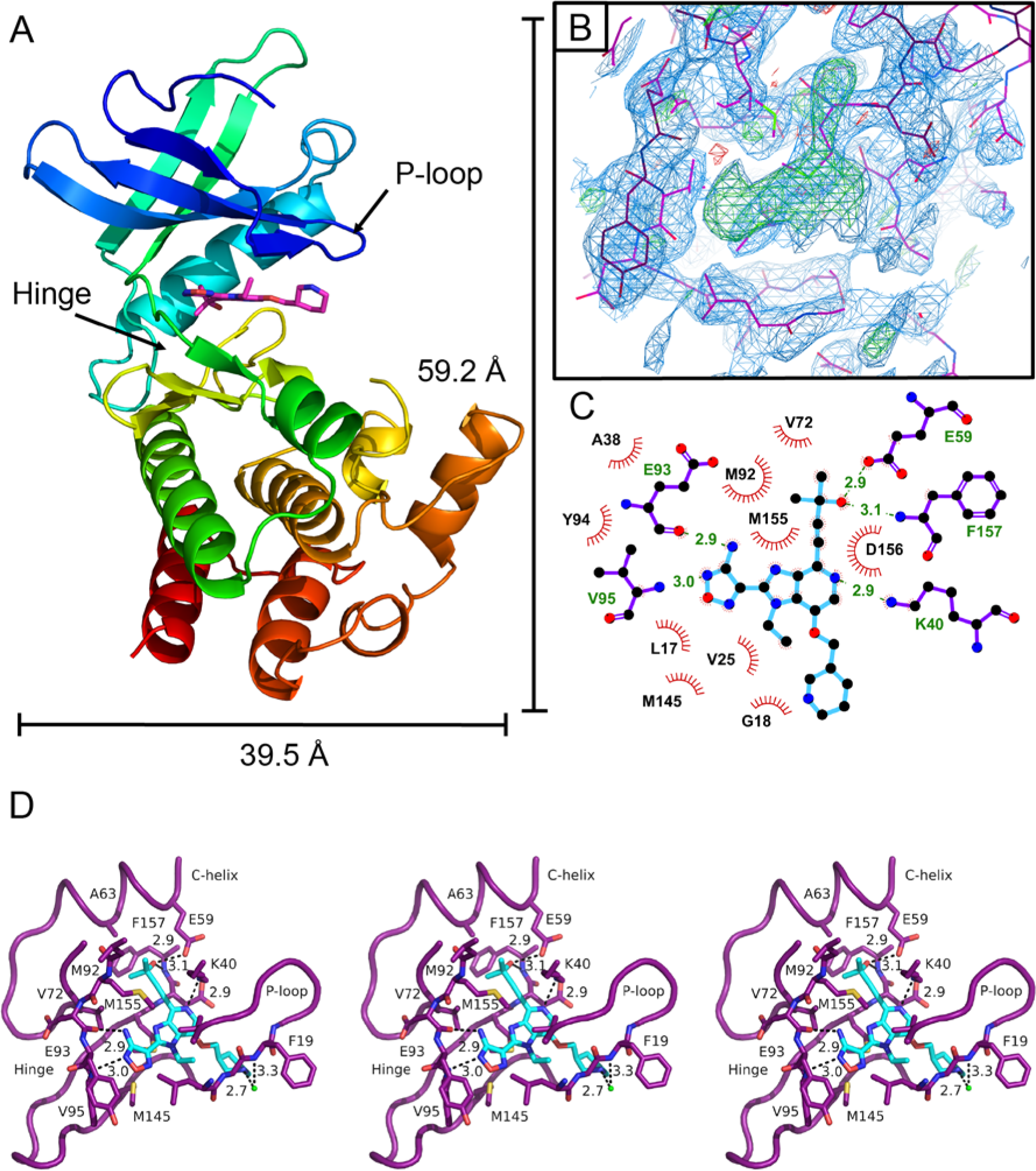
Structure of the PknB-GSK690693 co-crystal. **A)** The overall kinase structure is nearly identical to previous published structures (1O6Y, 2FUM, 1MRU) with overall dimensions as indicated. The hinge region and P-loop are also indicated. PknB is shown in N-C spectrum and GSK690693 is shown in magenta stick. **B)** The difference map in electron density after molecular replacement indicates the location of GSK690693 in the active site (green unmodeled density, 3σ). **C)** A ligand plot of GSK690693 interactions with PknB highlights several interactions. Hydrophobic residues are labeled in black with red hashes corresponding to red hashes around atoms in the ligand responsible for the interaction. Hydrogen bonds and corresponding donor or acceptor residues are labeled in green with distances in angstroms indicated. Hydrogen bonding residues are shown in purple sticks and GSK690693 is shown in cyan sticks. **D)** A 3D stereo image of the interactions between GSK690693 and PknB. Relaxed-eye stereo uses the left two images and cross-eye stereo uses the right two images. The P-loop should appear above the ligand. Hydrogen bonds are indicated with black dashes and the distance in angstroms is given. Residues participating in hydrogen bonding or stacking interactions are labeled. V72 and A63 are also labeled. Figures were rendered using Pymol (**A & D**), Coot (**B**), and Ligplot+ (**C**).

The position of GSK690693 in the PknB co-crystal structure corresponds to the most favored position calculated by Autodock (Fig. 4), with the imidazopyridine-aminofurazan core and the dimethylpropargyl alcohol in nearly identical positions in both the predicted and crystal structures. The model placed the R_2_ piperidine group inside the P-loop, whereas in the crystal structure it adopts a more extended conformation; however, this area is likely highly dynamic in solution. Overall, these data demonstrate GSK690693 binding in an ATP-competitive fashion, in agreement with our *in silico* modeling and previous structural biology for GSK690693 in Akt2.

**Figure 4.**
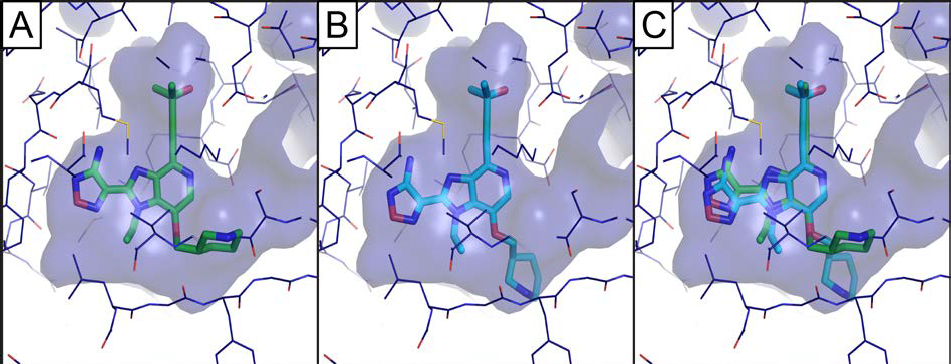
Docking poses agree well with the crystallographic model. **A)** The most energetically favorable position of GSK690693 (green sticks) in PknB (blue lines) as determined by Autodock. **B)** GSK690693 (cyan sticks) as modelled in PknB (blue lines) based on crystallographic data. **C)** The overlay of the docked pose and the actual position in the co-crystal shows generally good agreement. The active site cavity is outlined in transparent blue surface in all panels. The R2 group shows more variability in the docking models and is less well resolved in the crystal structure.

### Adding steric bulk in the PknB back pocket reduces inhibition by GSK690693

As most of the key hydrogen bond interactions in the back pocket are coordinated by backbone atoms or critical residues, and the numerous stacking and hydrophobic interaction residues individually contribute only a small fraction to binding, we focused on inhibiting entry to the back pocket to validate our SAR. We chose mutations V72M to restrict the back pocket entrance and A63I to sterically occupy the back pocket. The mutant enzymes were kinetically equivalent to the WT enzyme (**Fig. S5**), allowing us to use biochemical inhibition as a method to compare GSK690693 binding. Although the V72M mutant was statistically equivalent to wild type enzyme with GSK690693 IC_50_s of 483nM and 341nM respectively, occupying the back pocket was significantly more disruptive, as the IC_50_ for the A63I mutant was approximately 10-fold higher (IC_50_ = 3679nM) (Fig. 5). These results, combined with the ineffectiveness of SB747651A, support our hypothesis that binding the back pocket is critical for PknB inhibition by IPAs.

**Figure 5.**
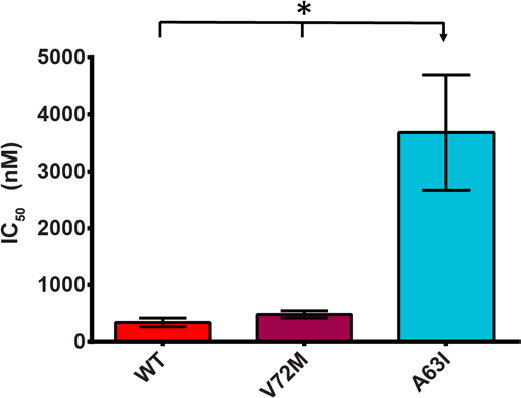
The back pocket mutant, A63I, is not efficiently inhibited by GSK690693. A63I mutant PknB has a higher IC_50_ than WT or a V72M (back pocket entrance) mutant. The difference in IC_50_ is statistically significant at p< 0.05 (indicated by *). The three mutants did not have statistically different K_m_ values (**Fig. S5**), indicating the change in inhibition is not due to changes in enzymatic activity.

### IPAs inhibit bacterial growth and can be potentiated by β-lactams

Effective biochemical inhibitors of PknB exist, but efficacy against whole mycobacteria has stagnated in the low micromolar range(Chapman et al., 2012, Lougheed et al., 2011). We initially tested *M. smegmatis* (*Msmeg*) as a safe and comparable surrogate for mycobacterial drugs in early development stages(Andries, Verhasselt et al., 2005). IPAs alone did not inhibit growth at 100μM or below, except for GSK1007102B which had an IC_50_ of 99μM. A sub-lethal dose of meropenem potentiated five out of eight IPA compounds to inhibit *Msmeg* growth, except for SB747651A, GSK554170A and GSK614526A (Fig. 6A), while GSK1007102B was the most potent inhibitor in combination. Importantly, IPA potentiation was specific to β-lactams as sub-lethal doses of isoniazid did not potentiate their microbiologic activity (**Fig. S7**), consistent with previous reports that that isoniazid and ethambutol did not improve the microbiologic activity of several structurally diverse, biochemically effective PknB inhibitors(Lougheed et al., 2011).

**Figure 6.**
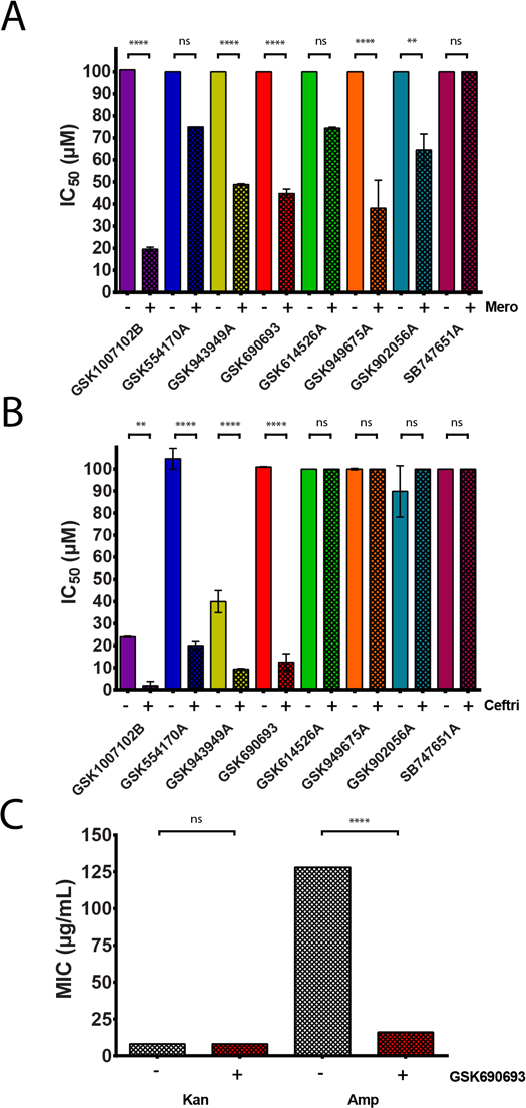
IPA compounds are potentiated by β-lactams in *M. smegmatis*, *N. asteroides*, and *M.abscessus* in culture. **A)** *Msmeg* and **B)** *Nast* were grown in various concentrations of IPA compounds and IC_50_ values were calculated (See **Figs. S7 & S9** respectively). **C)** Mab was grown with ampicillin and kanamycin both with and without GSK690693 and MICs were determined (**Table S3**). One way ANOVA was performed and statistically different groupings are denoted by ticks with arrows indicating the compound from which the group differs. Statistical significance is given by p < 0.05 (*), p < 0.01 (**), p < 0.001 (***), and p < 0.0001 (****). Statistics were calculated using PRISM (GraphPad), and the Tukey correction was used for multiple comparisons. MICs or IC_50_ values with β-lactam are given in **Fig. 1**, and compound colors correspond to those in **Fig. 1**. Hatched bars indicate β-lactam is present and solid bars represent drug only. All three species show significant potentiation by β-lactams relative to kinase inhibitor activity only.

The catalytic domains of PASTA kinases in the actinobacteria and firmicutes are highly conserved (**Fig. S8**). To determine whether this strategy is broadly applicable, we treated *Nocardia. asteroides* (*Nast*) with our IPAs. GSK1007102B, GSK554170A, GSK943949A, and GSK690693 were microbiologically effective with IC_50_ values of 24-100 μM, while the others had no measurable activity (Fig. 1C). Similar to *Msmeg*, these four inhibitors were significantly potentiated by β-lactams 4-10 fold relative to IPAs alone (Fig. 6B). These microbiologic IC_50_ values were only 5-30 fold higher than the biochemical IC_50_ values, indicating that β-lactam potentiation allowed some inhibitors to approach their maximum effectiveness. Taken together, the data suggest that IPAs could be broadly applicable across several PASTA kinase containing pathogens and that potentiation by β-lactams offers a significant improvement over IPA administration alone.

Although PknB is well conserved among mycobacteria (**Fig. S8**), differences in cell wall composition, efflux, and metabolism may complicate inhibitor accumulation in pathogens versus non-pathogens. We wanted to confirm whether IPAs would be effective at potentiating a β-lactam against a mycobacterial pathogen. We assayed the effect of GSK690693 against *M. abscessus* (*Mab*) both with a β-lactam and kanamycin, a non-β-lactam third line treatment option for resistant mycobacterial infections(Caminero, Sotgiu et al., 2010). Our results show that GSK690693 has no effect potentiating kanamycin; however, it did significantly potentiate the activity of ampicillin 8-fold (Fig. 6C). This agrees with the data in *Msmeg* that suggests this potentiation is specific to β-lactams and the 8-fold increase in effectiveness is similar to that seen in both *Msmeg* and *Nast*. Collectively our microbiological data point to the IPAs as authentic mycobacterial and nocardial inhibitors with the ability to be specifically potentiated by β-lactam antibiotics, forming a novel combination therapy.

## Discussion

Our data demonstrate that for well-defined targets, such as Hank’s-like kinases, *in silico* screening offers a cost-effective way to prioritize compounds for pharmacologic testing. We identified a family of inhibitors, the IPAs, that are biochemically active and bind the target, PknB, as our model predicted. These inhibitors were also microbiologically active and can be potentiated by normally ineffective β-lactams to form a novel co-therapy.

The computational model was highly predictive of the actual crystal structure (Fig. 4). The core scaffold, aminopyrazine ring, and dimethylpropargyl alcohol were in almost identical positions with similar hydrogen bonding patterns. The dimethylpropargyl alcohol at R_1_ was necessary for both biochemical and microbiological activity (Fig. 1C) suggesting that R_1_ position hydrogen-bonding in the back pocket is necessary for activity. Our structure shows two critical hydrogen bonds to the backbone and to E59, highlighting this importance. Although similar to the binding of GSK690693 in Akt2 (**Fig. S4**), multiple differences could be exploited to increase Akt-vs-PknB selectivity. The entrance to the back pocket is wider for PknB, and deeper on the E59 side than Akt (**Fig. S4**). Previous SAR with Akt shows that wider R_1_ groups (such as a phenyl) substantially inhibit an IPA’s activity(Heerding et al., 2008), indicating that this strategy is viable. Although most interactions between GSK690693 and PknB involve backbone hydrogen bonds or conserved residues (**Fig. S8**), our data suggest that point mutations that collapse the back pocket may be a possible resistance mechanism (Fig. 5). The back pocket is well conserved (**Fig. S8**), but to anticipate future resistance, the development of tolerated *in vivo* back pocket mutations should be studied. Although the computational model was less predictive of the R_2_ position, the basic orientation was correct, and based on the crystal structure, opportunities to improve the biochemical potency and selectivity exist.

We hypothesized that selecting small and hydrophilic inhibitors (Lipinski’s rules) would facilitate both entry into bacterial cells and future drug development efforts(Lipinski et al., 1997). Although effective antibiotics vary greatly in these parameters, most bacterial protein targeting drugs obey these guidelines(Mugumbate & Overington, 2015). Our IPAs could substantially inhibit growth of a non-pathogenic mycobacterium (*Msmeg*), a pathogenic mycobacterium (*Mab*), and a pathogenic non-mycobacterial actinobacteria (*Nast*) at micromolar levels when potentiated by a β-lactam (Fig. 1C). While other PknB inhibitors have been optimized with medicinal chemistry to achieve microbiological efficacy comparable to our IPAs(Chapman et al., 2012, Lougheed et al., 2011, Naqvi et al., 2014), our inhibitors have not, suggesting that sub-micromolar level inhibition may be attainable. While combination therapy is traditionally used for mycobacterial infections, the limited microbiologic activity of previously optimized PknB inhibitors in combination with traditional first line drugs such as isoniazid and ethambutol did not show any potentiation of activity(Lougheed et al., 2011). Previous data from our lab and others demonstrated that genetic deletion or pharmacologic inhibition of PASTA kinases in *Listeria monocytogenes* and *S. aureus* increases susceptibility to β-lactams(Pensinger et al., 2014, Vornhagen et al., 2015). Despite β-lactams’ general ineffectiveness against mycobacteria(Sotgiu et al., 2016) our decision to pair IPAs with β-lactams based on our previous data in firmicutes demonstrated significant benefit (Fig. 6). Additionally, although *Msmeg* and *Nast* did not require a β-lactamase inhibitor to see this effect, our *Mab* strain was a β-lactamase knockout; therefore, a tripartite therapy with a β-lactamase inhibitor may be required to protect the β-lactam in organisms with a different β-lactamase profile. Although we cannot rule out inhibition of other ten mycobacterial kinases (such as PknA and PknG) by IPAs, but the data are suggestive of growth inhibition being inhibited at least in part by PknB. Genetic data show that β-lactam potentiation is not seen in Listeria monocytogenes or MRSA harboring PASTA kinase knockouts. Additionally, IPAs definitively bind PknB (Fig. 3) with high affinity (Fig. 1 & 2), and PknB is the only mycobacterial serine/threonine kinase to have penicillin binding domains(Prisic & Husson, 2014), suggesting a route for β-lactam potentiation.

Furthermore, as the other mycobacterial kinases are also involved in growth and virulence(Prisic & Husson, 2014), off-target inhibition of these structurally similar kinases should only serve to enhance its drug potential.

Taken together, our data suggest that computational modelling accurately predicted the binding modes and biochemical activity of the IPA family of drugs. These un-optimized PknB inhibitors have mycobacterial activity similar to other discovered inhibitors, but less of a discrepancy between biochemical potency and microbiologic potency than known inhibitors, potentially suggesting better bacterial access/retention. Importantly, we found that combining a β-lactam with biochemically “average” PknB drugs can create microbiological inhibition near levels seen with other inhibitors, opening the field for further improvements. This approach allowed us to leverage a large body of industrial and academic work in a cost-efficient and timely manner to repurpose a failed drug as a new lead for a specific antibiotic to use in a novel co-therapy.

## Materials and Methods

### Docking

Coordinates for PknB (1O6Y) were downloaded from the protein data bank and prepared for docking using Sybyl 2.1.1(Tripos). Ligands and waters were removed and Gasteiger-Huckel charges were added. 3D coordinates for ligands in the PKIS libraries were obtained from the Small Molecule Screening Facility at UW-Madison(Drewry et al., 2014). The structure data file for the Selleck library was obtained from Selleck and converted into 3D coordinates using OMEGA (OpenEye Scientific). OMEGA was also used to add AM1BCC charges to the ligands in all libraries. A total of 1054 unique kinase inhibitors from three libraries (Selleck, PKIS 1, and PKIS 2) were prepared. Autodock Tools 1.5.6 was used to prepare the receptor coordinates (PknB) for docking(Morris, Huey et al., 2009). Docker 1.0.4 script was modified to retain supplied charges in the ligand files and was then used to automate ligand docking to the receptor using Autodock 4(Morris et al., 2009). Docking was done at the rate of 30 iterations per ligand using a Lamarckian genetic algorithm. Docking results were organized and graphed using Excel (Microsoft) and PRISM (GraphPad). Poses were viewed using Pymol (PyMOL Molecular Graphics System, Version 1.7.6.0 Schrödinger, LLC.) with the Autodock/Vina plugin(Trott, Department of Molecular Biology et al., 2016). Chemical structures for figures were rendered using ChemDraw (Perkin Elmer). Lipinski’s rules were calculated using Sybyl and used to further narrow down selection of our starting compounds(Lipinski et al., 1997).

### Chemicals and Reagents

All chemicals were purchased through Fisher Scientific unless otherwise noted. Miniprep kits and IPTG were from IBI scientific. Restriction enzymes were from New England Bio Labs. DTT and Ampicillin were from Dot Scientific. LB was from Growcells. GS4FF resin was from GE Healthcare. Meropenem was from APP Pharmaceuticals. Magnesium chloride, ATP·Mg, PMSF, chloramphenicol, glycerol, and bis-tris propane were from Sigma. PEG 3350 was from Microscopy Sciences. ATPγ_32_P was obtained from Perkin Elmer.Thymine was from Acros Organics. β-cyclodextran was from Alfa Aeser. GSK690693 was from Selleck Chemicals. Other IPA compounds were acquired through collaboration with co-authors at UNC and their synthesis is published (Heerding et al., 2008, Rouse et al., 2009).

### Cloning

PknB 1-331 and full length GarA were amplified from Mycobacterium tuberculosis genomic DNA using Phusion polymerase (ThermoFisher) and standard cycling conditions.Amplified DNA was digested and ligated into PGEX-6P vector (GE) and transformed into Top10 competent cells for plasmid storage and BL21-Rosetta cells for protein production. Sequencing was done at the UW BioTechnology Center to confirm the constructs. PknB 1-331 A63I and V72M mutants were generated by following the Quikchange Mutagenesis protocol (Agilent) with minor changes. Primers were designed using Agilent Quikchange Primer Design (IDT) and used in PCRs using PrimeStar polymerase (NEB). PCR product was exposed to EtOH precipitation overnight in 100% EtOH at −20C. The following day the DNA was collected at 16,060g and washed with 75% EtOH before being resuspended in water. The DNA was then digested with DPNI overnight at 16C. The digest was directly transformed into Top10 cells and plated on LB agar containing 1mM ampicillin. Colonies were picked and inoculated overnight at 37C, and DNA was extracted and sequenced. The plasmid with the correct sequence was transformed into BL21-Rosetta cells for protein expression. Codon optimized PknB 1-280 cDNA was ordered as a gBlock (IDT) and expression constructs were prepared as described above.

### Protein Purification

Initial affinity purification of all PknB constructs was performed using the following procedure. LB broth containing 1mM ampicillin and 1mM chloramphenicol was inoculated with overnight cultures at 37C. Cultures were induced with 0.5mM IPTG after reaching an OD_600_ of 0.6-0.8. Protein was overexpressed overnight at 23C. Cells were harvested at 6,079g and resuspended in lysis buffer [25mM Tris pH 8.0, 150mM NaCl, 1mM DTT, and 10mM MgCl_2_] with 1mM PMSF and DNAse. The lysate was sonicated and the soluble fraction was collected at 30,600g and passed over columns with GS4FF resin (GE) at approximately 4 mL of resin per liter of culture. Columns were rinsed with lysis buffer, and protein was removed from the resin with elution buffer [50mM Tris pH 8.0, 5mM NaCl, 10mM MgCl_2_, 3mM DTT, and 20mM reduced glutathione (GSH)] and digested with 1/20 w/w PreScission protease. GarA was produced using the same procedure except MgCl_2_ was omitted from the buffers.

Digested PknB 1-331 WT and mutants were separated from GST and GSH using anion exchange (Source 15Q, GE) using buffer A [20 mM Tris pH 8.0, 1 mM DTT, 1mM MgCl2] and buffer B [buffer A + 1M NaCl] in a stepped gradient as follows: 0-20 % B for 0.5 column volumes (CV), 20-22.5% B for 5 CV, and 22.5-50% B for 5 CV. PknB was concentrated using 10kDa cutoff columns (Millipore) at 3150g and injected onto a Superdex 75 column (GE) for size exclusion purification using a buffer containing 20 mM Tris pH 8.0, 1 mM DTT, 1mM MgCl2, and 150mM NaCl. Protein was collected at 95% or greater purity and was flash frozen for storage at -80C.

Digested GarA was partially separated from GST using anion exchange with a gradient of 0-19% B for 0.5 CV and 19-50% B for 5 CV. GarA was passed over GS4FF resin (∼1mL resin per 6mg protein) three times to remove GST before concentration and size exclusion. Protein was collected at 95% or greater purity and was flash frozen for storage at -80C. All buffers were the same as for PknB except MgCl_2_ was omitted.

Digested PknB 1-280 for crystallization was further purified using the above procedure, except after anion exchange, the protein was passed over GS4 FF resin (∼1mL resin per 6mg protein) to remove traces of GST before concentration and size exclusion. After size exclusion, PknB was concentrated to ∼10mg/mL, a small sample was saved to measure activity, and GSK690693 was added to the protein at 3x the molar concentration (330μM) using the following procedure. Two mL of buffer from size exclusion was warmed to 37C and 612μL of 1mM GSK690693 in DMSO was added slowly added to the warm buffer with frequent mixing. This mixture was added to 2.5mL of PknB and allowed to incubate at room temperature for 15min.After incubation, the sample was spun for 5min at 3150g to remove precipitate then concentrated to 9.8mg/mL. Kinase inhibition was checked using the KinaseGlo® assay as described in the “Kinase assays” section.

### Kinase assays

The kinase assays were performed using the KinaseGlo® reagent from Promega. All reactions were done in 50μL volume. The buffer used for all kinase assays was 10mM Tris-HCl pH 7.4, 150mM NaCl, 1mM DTT, and 1mM MgCl_2_. For kinetic analysis, plates were prepared with a serially diluted concentration gradient of GarA from 0-104μM across a serially diluted gradient of ATP from 0 to 400μM such that each concentration of GarA was present at each concentration of ATP. 0.25μM of PknB was added to each well and allowed to react at 37C for 20min, then equilibrated to room temperature for 10min. After incubation, the reaction was stopped by the addition of 50μL of KinaseGlo® reagent, and the signal was allowed to stabilize for 10min at room temperature per the product manual. The plate was read using luminescence detection on a Synergy HT detector (BioTek) and the data were collected using the Gen5 2.0 software (BioTek). The data were processed in Excel (Microsoft) and non-linear regression models were fit in PRISM (GraphPad). The model for the velocity of the reaction plotted as a function of GarA concentration was fit using an allosteric sigmoidal model in preference over a Michaelis-Menten model (p <0.0001) with a shared Hill coefficient (**Fig. S1A**). The model for velocities as a function of ATP concentration was fit using the Michaelis-Menten model over the allosteric sigmoidal model (p <0.0001) (**Fig. S1B**). Curves were fit to plotted apparent V_max_ and true V_max_ values were calculated based on the upper limit of each curve (**Fig. S1C & D**). K_cat_ was calculated based on Vmax using the equation V_max_ = K_cat_ * [E_t_] where [E_t_] is the total enzyme concentration (0.25μM). True K_m_ values for ATP and GarA were calculated similarly to true V_max_ values (**Fig.S1E & F**).

Inhibition kinetics were determined using similar parameters as for enzyme kinetics with the following changes. Drugs in 5mM DMSO were diluted in kinase buffer to 3/5 the final concentration using a serial dilution from 20 or 10μM to 0μM. The final DMSO concentration in the reactions was no more than 0.4%. PknB 1-331 was added to the drugs to a final concentration of 0.25μM and allowed to incubate for 10min at 37C. ATP and GarA were added for a final concentration of 100μM and 42μM respectively, initializing the reaction. The reaction proceeded and was quantified as described for enzyme kinetics. The data were transformed to log scale and non-linear regression was performed in PRISM using the variable slope 4-parameter model for enzyme inhibition to determine IC_50_ (**Fig. S2**). K_i_ values were determined using the following equation(Heerding et al., 2008):

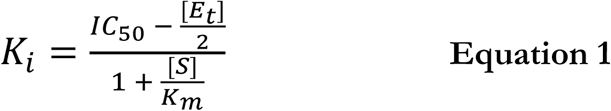

where [E_t_] is the total enzyme concentration (0.25μM), [S] is the total substrate (ATP) concentration (100μM), and K_m_ is the K_m_ for ATP (42μM).

Radiolabeled kinase assays were done as follows. The reaction buffer was the same as above and reactions were performed in 20μL volumes. PknB was used at a final concentration of 0.5μM, cold ATP was 20μM, ATPγ_32_P was 0.1nM (10μCi), GarA was 10μM, and GSK690693 was serially diluted from 0-20μM. GSK690693 and PknB were incubated together for 10min at 37C then substrates were added and allowed to react for 1hr at 37C. The reaction was stopped by the addition of 10μL SDS loading buffer and 20μL of the resulting mixture was loaded on a 15% gel for SDS-PAGE. The gel was fixed with a solution of 10% acetic acid, 40% methanol, and 5% glycerol for 1hr. The gel was dried for 1hr and autoradiography film (MidSci) was exposed to the gel for 2min.

### Crystallization and structure determination

PknB 1-280 for crystallization was produced as described above. Protein was sent to the Hauptman-Woodward institute for screening in 1536 different crystallization conditions in the microbatch-under-oil format at room temperature(Luft, Collins et al., 2003). Hits were visually assessed using the Macroscope J program (Hauptman-Woodward). The initial hit condition was 0.5 M bis-tris propane pH 6.7 and 15% w/v PEG 3350. This was optimized with a matrix of PEG and buffer concentrations and additives to grow larger, single crystals. Final diffracting crystals were produced as follows: 0.5μL of buffer A [25% PEG 3350, 0.5M bis-tris propane pH 7.5, 1% glycerol v/v] was combined with 0.5μL buffer B [0.2% thymidine w/v and 0.2% β-cyclodextran w/v] were combined with 1.0μL PknB (9.8 mg/mL in size exclusion buffer listed above). The drops were equilibrated at room temperature in the sitting drop format over 50μL of buffer A. Crystal growth was monitored over the course of two weeks and crystals which showed no visible growth in consecutive viewings were harvested and cryopreserved. Crystals were cryopreserved with the following procedure: crystals were resuspended in well buffer containing 0.3mM GSK690693 and slowly transferred to a cryobuffer containing 30% glycerol, well buffer, and 0.3mM GSK690693 in a 1:1 ratio. The solution was removed and the procedure was repeated five times until the final concentration of the solution surrounding the crystal was essentially the same as the cryobuffer. Crystals were mounted in loops (Molecular Dimensions) and quickly flash frozen in liquid nitrogen.

Diffraction data was collected at Sector 21, beamline D at the Advanced Photon Source (APS) at Argonne National Laboratory (ANL) and processed with HKL2000(Otwinowski & Minor, 1997). Model building and refinement was done using the PHENIX software package (Adams, Afonine et al., 2010). Data were assessed using Xtriage, and molecular replacement was done using Phaser-MR with 1O6Y as the search model. Ligand coordinates were generated with eLBOW, and the structure was refined with phenix.refine. The model was manually built and adjusted in Coot and further refined with phenix.refine(Adams et al., 2010, Emsley & Cowtan,2004). Coordinates were deposited in the Protein Data Bank under accession code 5U94. Figures were created using Pymol, Coot, and Ligplot+(Emsley & Cowtan, 2004, Wallace, Laskowski et al., 1995).

### Cell Viability Assays

*Mycobacterium smegmatis* (*Msmeg*) was streaked out onto lysogeny broth (LB) agarose plates from a frozen stock. After incubation at 37°C for 3 days, a colony was picked and inoculated into a 3mL culture of Middlebrook 7H9 media supplemented with 0.08% tyloxapol (v/v) at 37°C and grown overnight. Culture density was measured and then diluted to an OD_600_ reading of 0.3. IPA compounds were serially diluted as appropriate in a 96-well plate. If the addition of meropenem was being tested, 40μL of meropenem (at concentration of 3.125μg/mL) was added to the serially diluted 50μL of IPA compound. If no meropenem was being tested, 40μL of Middlebrook 7H9 media supplemented with 0.08% tyloxapol was added instead. 10μL of *Msmeg* at OD_600_ of 0.3 was added to the previous mixture to give a final total volume of 100μL. The final concentration of meropenem (if added) was 1.25μg/mL and the final concentration of IPA ranged from 0-100μM. 100μL of Middlebrook 7H9 media with 0.08% tyloxapol (v/v) was used as a negative control.10μL of *Msmeg* added to 90μL of Middlebrook 7H9 with 0.08% tyloxapol (v/v) was used as a positive control. The plate was incubated for 16 hours at 37°C before 10μL of resazurin (1mg/mL) was added to each well. After four hours of incubation at 37°C, resazurin fluorescence was read on a microplate reader using an emission wavelength of 530nm and an excitation wavelength of 590nm. The data obtained was then converted into % growth by comparing the control wells (bacteria alone) with the experimental wells. The data were graphed in PRISM (GraphPad) and nonlinear regression was used to calculate IC_50_ values using the same methods as for enzyme inhibition. ANOVA was performed using PRISM (GraphPad) using the Tukey correction for multiple comparisons. Ineffective inhibitors were analyzed at concentration of 100μM, representing the minimum difference possible based on our assay conditions.

The same was done with *Nocardia asteroides* (*Nast*) with minor changes. Nast was grown from frozen stock into LB where it grew overnight. The following day, Nast was diluted to an OD600 of 0.3. Additionally, ceftriaxone was added in place of meropenem, with the final concentration of ceftriaxone being 0.25μg/mL. The 96-well plate was only incubated for 16 hours at 37°C before 10μL of resazurin (1mg/mL) was added to each well. The resazurin was incubated with Nast for 4 hours at 37°C before being read on the microplate reader.

*Mycobacterium abscessus* PM3386 δbla (*Mab*) MIC determinations were done according to the microdilution method. Antibiotics were tested in two-fold serial dilutions with GSK690693 at 50 uM or with a DMSO control in a 96-well format. Other control wells with cells received no antibiotics. The strain was streaked out onto Middlebrook 7H9/ADS agarose plates from a frozen stock. After incubation at 37°C for four days, an *Mab* colony was picked and inoculated into 10 mL of Middlebrook 7H9 ADS media supplemented with 0.5% tyloxapol (v/v) and grown with shaking at 37°C. The next day, the culture was adjusted to an optical density of 0.5 (OD_600)_), diluted 100-fold, and 100 μl of the suspension used to inoculate each well. The plates were incubated for 48 hours, stationary, at 37°C, and then 50 υL of 1:1 mixture of AlamarBlue:10%(v/v) Tween-80 was added to each well and the plate incubated for an additional 24 hours at 37°C. The wells that remain blue were scored as the MIC (μg/mL).

## Acknowledgments

We would like to thank Steve Darnell, Spencer Erikson, and Scott Wildman from the Small Molecule Screening and Selection Facility for their assistance with scripts for docking and ligand preparation. We would also like to thank Caitlyn Pepperell for providing *Mycobaterium tuberculosis* genomic DNA and Kyle Boldon and Rehan Tariq for help with cloning, growth assays, and protein purification. We would particularly like to thank Andrew Mehle for allowing us to use his ÄKTApurifier. Additionally, we would like to thank Jim Keck for allowing us to use his in-house X-ray equipment and David Smith at the APS for beamline assistance and setup. This work was supported by grant UL1TR000427 from the Clinical and Translational Science Award (CTSA) program of the National Center for Advancing Translational Sciences, NIH, the Hartwell Foundation, NIAID/NIH/HHS grant AI500145, NIH/NCI P30 and CA014520.

## Author Contributions

N.W. oversaw the project, designed the experiments, performed all virtual docking, biochemistry, and structural biology experiments, analyzed the data, rendered the figures, and wrote the paper. N.T. did technical development for microbiologic assays and collected *M. smegmatis* data.R.P. did cloning, protein purification, and phylogeny analysis. J.B. collected *N. asteroides* data. A.S. assisted with microbiologic assays and cloning. K.S. provided modeling consultation, M.P. performed *M. abscessus* assays and provided the associated data. B.Z. and D.D. provided all IPAs except GSK690693, edited the manuscript, and provided insight into kinase inhibitor libraries. JD.S. provided assistance with experimental design and edited the manuscript. R.S. conceived and oversaw the project, edited the manuscript, and provided direction for technical problems, writing, and submission.

